# The Ufm1 system is upregulated by ER stress during osteogenic differentiation

**DOI:** 10.1101/720490

**Authors:** Michal Dudek, Gillian A. Wallis

**Affiliations:** Wellcome Centre for Cell Matrix Research, Division of Cell Matrix Biology and Regenerative Medicine, School of Biological Sciences, Faculty of Biology, Medicine & Health, Manchester Academic Health Science Centre, University of Manchester, Oxford Road, Manchester M13 9PT, UK

**Keywords:** Ufm1, Ufsp2, ER Stress, Osteoarthritis

## Abstract

**Objective:** Mutations in the catalytic site of the ubiquitin-fold modifier 1(UFM1)-specific peptidase 2 (UFSP2) gene have been identified to cause autosomal dominant Beukes hip dysplasia in a large multigenerational family and a novel form of autosomal dominant spondyloepimetaphyseal dysplasia in a second family. We investigated the expression of the UFSP2/UFM1 system during mouse joint development the connection to ER stress induced during osteogenic differentiation.

**Methods:** The pattern of expression of *Ufsp2* was determined by radioactive RNA in situ hybridisation on mouse tissue sections. qPCR was used to monitor expression during *in vitro* osteogenic differentiation and chemically induced ER stress. Affinity purification and mass spectrometry was used for isolation and identification of Ufm1 conjugation targets. Luciferase reporter assay was used to investigate the activity of Ufm1 system genes’ promoters.

**Results:** We found that *Ufsp2* was predominantly expressed in the bone and secondary ossification centres of 10-day old mice. The Ufm1 system was upregulated during *in vitro* osteogenic differentiation and in response to chemically induced ER stress. We identified unfolded protein response elements in the upstream sequences of *Uba5, Ufl1, Ufm1*and *Lzap*. We identified putative Ufm1 conjugation targets where conjugation was increased in response to ER stress.

**Conclusion:** Higher expression of Ufsp2 in bone and secondary ossification centres as well as upregulation of components of the Ufm1system in response to ER stress suggests that the molecular pathway between the *UFSP2* mutations and form of skeletal dysplasia may relate to abnormal ER stress responses during osteoblast differentiation.

## 1. Introduction

We have previously identified a unique mutation in the Ufm1-specific peptidase 2 gene, *UFSP2*, that results in loss of UFSP2 catalytic activity (*in vitro*) of in a large multigeneration family with Beukes Hip Dysplasia (BHD) [1]. BHD is an autosomal dominant disorder characterised by bilateral dysmorphism that is limited to the proximal femur and delayed ossification of the secondary centre, which results in severe progressive degenerative osteoarthropathy from childhood. A second novel mutation in the *UFSP2* gene, also predicted to cause loss of UFSP2 catalytic activity [2] has been identified in affected members of a family with a novel form of autosomal dominant spondyloepimetaphyseal dysplasia (SEMD) involving the epiphyses predominantly of the hips, but also of knees, ankles, wrists and hands, with variable degrees of metaphysis and spine involvement and characterised by short stature, joint pain and genu varum.

UFSP2 is a component of the Ufm1 (ubiquitin-fold modifier 1) system, which like the ubiquitin system, comprises a multi-enzyme cascade the major components of which are: i) the E1-activating enzyme, Uba5 (ubiquitin like modifier activating enzyme 5); ii) the E2 enzyme, Ufc1 (UFM1 conjugating enzyme 1), responsible for conjugation of Ufm1 to target proteins; iii) Ufl1, a Ufm1 E3 ligase, and iv) Ufsp1 and Ufsp2 (the UFM1 specific peptidases 1 and 2) [3]. Additionally, Lzap (Cdk5rap3) is a substrate adaptor for Ufl1 and Ddrgk1 is one of few known Ufm1 conjugation targets. Loss-of-function mutation in *DDRGK1*gene was found to cause Shohat-type SEMD, a condition characterised by disproportionate short stature, lordosis, genu varum, and joint hypermobility. Radiographically, patients with SEMD have delayed epiphyseal ossification, platyspondyly with central notches in the vertebral end plates, radiolucency of the femoral metaphyses, and relative fibular overgrowth. Genetic deletion of *DDRGK1*in zebrafish and in mice led to disruption of cartilage formation by decreasing Sox9 protein level [4].

Components of the Ufm1 system localise to the ER, and functional studies point to specialised roles for ufmylation in regulating ER-related stress [3]. The ER is a major site of protein quality control, housing the synthesis machinery for proteins destined to be secreted or membrane-trafficked within the cell, as well as chaperones to assist protein folding. Chondrocytes and osteoblasts actively secrete the proteins that form the vast extracellular matrix of cartilage and bone. Forms of skeletal dysplasia caused by mutations in matrix genes have been identified where ER stress is a common feature of their molecular pathology [5, 6].

In this study, following identifying that Ufsp2 is expressed by osteoblasts during mouse joint development, we investigated whether the Ufm1 system is differentially regulated during osteogenic differentiation and in response to ER stress.

## 2. Methods

### 2.1 Radioactive in situ hybridisation

#### Probe synthesis

DNA template for the synthesis of RNA probe was prepared by linearisation of plasmids containing probe sequence with a restriction enzyme cutting at the 5’ end of RNA polymerase transcription site generating antisense RNA product. The template DNA was then separated on agarose gel and extracted using QIAquick Gel Extraction Kit. The radio-labelled probe was synthesised replacing UTP with ^35^S UTP using Riboprobe® System (Promega) for 40 min at 37°C. After this time 1μl of RNA polymerase was added and the reaction was incubated at 37°C for an additional 1 hour. Following transcription the DNA template was digested with DNase at 37°C for 10 minutes. The RNA probe was then precipitated. The pellet was washed with 50 μl of 10 mM DTT diluted in 100% EtOH. After drying the RNA pellet was resuspended in 50 μl of 50 mM DTT diluted in DEPC H2O.

#### Preparation of tissue sections

Mouse tissues were fixed overnight in 4% PFA in PBS, processed in Shandon Citadel 2000 tissue processor and paraffin embedded. Tissues from mice older than newborn were decalcified for three days in decalcification solution (20% EDTA, 4% PFA in PBS pH 7.4) at 4°C before embedding. Slides were prepared by sectioning paraffin blocks on Microm HM 355S microtome with feed set to 5 and drying them overnight in 50°C. The sections were then de-waxed in xylene and rehydrated by passing through a graded ethanol series. The slides were incubated in 2% HCl for 20 minutes, washed in 2x SSC and incubated in proteinase K buffer for 10 minutes at 37°C. The proteinase was then inactivated by dipping the slides in 2mg/ml glycine for 2 minutes at room temperature. Slides were washed twice in PBS, incubated for 20 minutes in 4% PFA-PBS and washed in PBS again. The slides were then incubated in triethanolamine (triethanolamine 4.65 ml, 875 μl acetic anhydride and 345ml H2O) for 10 minutes at room temperature and washed in PBS followed by wash in H2O and dehydrating through the graded ethanol series.

#### Hybridisation

The probe was measured using a scintillation counter and the amount of probe to be used per slide was calculated according to the following formula: The required amount of probe was added to the hybridisation buffer (100 μl of buffer per slide) and heated to 95°C. The hybridisation mix was then applied on the slides and the slides were covered with glass cover slips. The slides were then placed in hybrydisation chamber containing 50% formamide and incubated overnight at 55°C. On the next day cover slips were removed and slides were incubated in following wash solutions and buffers: Wash solution twice at 55°C for 20 min and once at 65°C for 20 minutes; RNase buffer twice at 37°C for 15 minutes; RNase A at 37°C for 30 minutes; RNase buffer at 37°C for 15 minutes; Wash solution twice at 65°C for 20 min; SSC/DTT buffer at 5°C for 20 min and then 5 minutes at room temperature; ammonium acetate / ethanol for 2 minutes at room temperature. The slides were then dehydrated through the graded ethanol series, air dried and exposed to autoradiography film overnight to confirm the hybridisation worked before covering the slides with photoreactive emulsion.

#### Developing

After confirming the hybridisation with autoradiography film the slides were covered with K5 nuclear emulsion (Ilford, UK) prepared as 50:50 solution with 2% glycerol, dried and stored in dark for 2 weeks before developing. The slides were developed in D19 developer (Kodak) and fixed using Unifix (Kodak). After developing slides were H&E stained using Thermo Shandon Linistain GLX stainer.

### 2.2 Protein extraction

Cartilage tissues were pulverized using a liquid-nitrogen-cooled tissue grinder and proteins extracted as previously described [5]. Briefly, cartilage samples were reconstituted in 100 μL of 100 mM Tris acetate buffer pH 8.0 containing 10 mM EDTA and protease/phosphatase inhibitors and deglycosylated by treatment with 0.1 units of chondroitinase ABC for 6 h at 37 °C. Proteins were sequentially extracted in a chaotropic buffer containing guanidine hydrochloride (4M GuHCl, 65 mM DTT, 10 mM EDTA in 50 mM sodium acetate, pH 5.8). Protein samples were precipitated with nine volumes of ethanol, washed once in 70% ethanol, then resuspended in 120 μL of solubilisation buffer (7 M urea, 2 M thiourea, and 30 mM Tris, pH 8.0) and the volume was adjusted to achieve a concentration of ∼1 mg/mL, as estimated using the EZQ protein quantitation protocol (Thermo Fisher). Samples were then stored at −80 °C until required. Protein samples were analysed by SDS-PAGE and detected by silver staining as previously described.

### 2.3 Osteogenic differentiation

Subconfluent 2T3 cells were trypsinized and plated on gelatine coated 12 well plates at a density of 1.6×104 cells/cm^2^. On reaching confluency (usually day 3) the cells were cultured in MEM (alpha modification) containing 5% FBS (v/v), 2 mM L-glutamine, 1x penicillin/streptomycin, 100 μg/ml L-ascorbic acid 2-phosphate, 5 mM β-glycerophosphate and 10 ng/ml rhBMP-2. Media was changed every 2 days. The cultures were usually maintained until day 12.

### 2.4 Luciferase assay

2T3 cells were plated in 96 well solid white microplate (Fisher, UK) at a density of 80% per well in 100 μl of growth medium. Cells were transfected using FuGene reagent in a 3:1 DNA to reagent ratio according to the manufacturer’s protocol. Each well was co-transfected with two plasmids: the pGL3-basic luciferase vector carrying 70 the tested genomic sequence and a control plasmid carrying *Renilla* luciferase gene under the control of the CMV promoter. The plasmids were co-transfected in 2:1 ration (ng/ng). All transfections were done in triplicate. 24h post transfection the medium was changed to fresh medium with or without 2μM thapsigargin. After 14h of incubation the media was removed and 25μl of PBS was added to each well followed by 25μl of Dual-Glo Luciferase Assay Reagent. The pGL3 firefly luciferase luminescence was measured after 10 minutes of incubation in a luminometer. Next, 25μl Dual-Glo Stop & Glo reagent was added, incubated for 10 minutes and *Renilla* luciferase luminescence was measured as previously. The values were calculated as ratio of firefly:*Renilla* luminescence for each well.

### 2.5 Affinity purification using His-Tag

#### Preparation of cell lysates

Culture media was removed and the cells were washed with ice cold PBS with protease inhibitors. PBS was removed and Gu-HCl denaturing lysis buffer (6M Guanidinium-HCl, 100 mM Na_2_HPO_4_/ NaH_2_PO_4_ buffer pH 8.0, 10 mM Tris-HCl pH 8.0, 10 mM imidazole, 10 mM β-mercaptoethanol, 20 mM N-ethylmaleimide, 20 mM iodoacetamide, 1x cOmplete protease inhibitors) was added (0.4 ml per well of a 6 well plate or 4 ml per T75 flask). The cell lysates were collected in to tubes and were allowed to lyse further for 1 hour at 4°C on a rotary shaker set to low rpm. After the incubation the lysate was passed 4 times through 21G, 23G, and 25G needles to reduce the viscosity. The lysate was then centrifuged at 4°C and 13000 rpm for 30 minutes and decanted to a new tube avoiding the carryover of the cell debris collected at the bottom of the tube.

#### Affinity purification

Ni-NTA resin (5Prime) was directly added to the cleared cell lysate. The resin was incubated with the lysate for 4h or overnight at 4°C on a rotary shaker set to low rpm. On the next day the resin was briefly centrifuged and the supernatant (flow-through) was kept for subsequent analysis. The resin was washed with lysis buffer and then with Urea buffer A (8M urea, 100 mM Na_2_HPO_4_/ NaH_2_PO_4_ buffer pH 8.0, 10 mM Tris-HCl pH 8.0, 10 mM imidazole) (twice) and Urea buffer B (8M urea

100 mM, Na_2_HPO_4_/ NaH_2_PO_4_ buffer, 10 mM Tris-HCl, 10 mM imidazole, pH6.3) (twice). The resin was washed with neutral Buffer W before eluting with 1 resin volume of Elution Buffer (50 mM NaH_2_PO_4_, 300 mM NaCl, 250 mM imidazole, pH7.0)

### 2.6 Mass Spectrometry

#### Digestion

Bands of interest were excised from the gel and dehydrated using acetonitrile followed by vacuum centrifugation. Dried gel pieces were reduced with 10 mM dithiothreitol and alkylated with 55 mM iodoacetamide. Gel pieces were then washed alternately with 25 mM ammonium bicarbonate followed by acetonitrile. This was repeated, and the gel pieces dried by vacuum centrifugation. Samples were digested with trypsin overnight at 37 °C. The samples were extracted in one wash of 20mM ammonium bicarbonate, and two of 50% acetonitrile, 5% formic acid. The extract was then dried by vacuum centrifuge to 20µl.

#### Mass Spectrometry

Digested samples were analysed by LC-MS/MS using an Ultimate 3000 (LC-Packings, Dionex, Amsterdam, The Netherlands) coupled to a HCT Ultra ion trap mass spectrometer (Bruker Daltonics, Bremen, Germany). Peptides were concentrated on a pre-column (5 mm × 300 μm i.d, LC-Packings). The peptides were then separated using a gradient from 98% A (0.1% FA in water) and 1% B (0.1% FA in acetonitrile) to50 % B, in 40 min at 200 nL min^−1^, using a C18 PepMap column (150 mm × 75 μm i.d, LC-Packings).

#### Data Analysis

Data produced was searched using Mascot (Matrix Science UK), against the UniProt database of all mouse proteins. Data were validated using Scaffold (Proteome Software, Portland, OR).

## 3. Results

### 3.1 Ufm1 system and osteogenic differentiation

Radioactive *in situ* RNA hybridisation was performed on tissue sections from E12.5, E14.5, new-born and 10 day old mice. No *Ufsp2* expression was detected in serial tissue sections of either embryonic or new-born mice (Supplementary Figure 1). In the hip region of 10 day old mice the *Ufsp2* expression pattern resembled that of *Col1a1. Ufsp2* expression was also visible in the muscle but not in the cartilage of the femoral head or the acetabulum (Figure 1A). In the knee, (Figure 1B) *Ufsp2* expression was detected in the bone and muscles, the secondary ossification centre and, with weaker expression in part of the proliferative zone of cartilage. *Ufsp2* expression was also detected in the spine (Figure 1C) where the signal was in the bone of the vertebral bodies, annulus fibrosus and the end plate.

**Figure 1.**
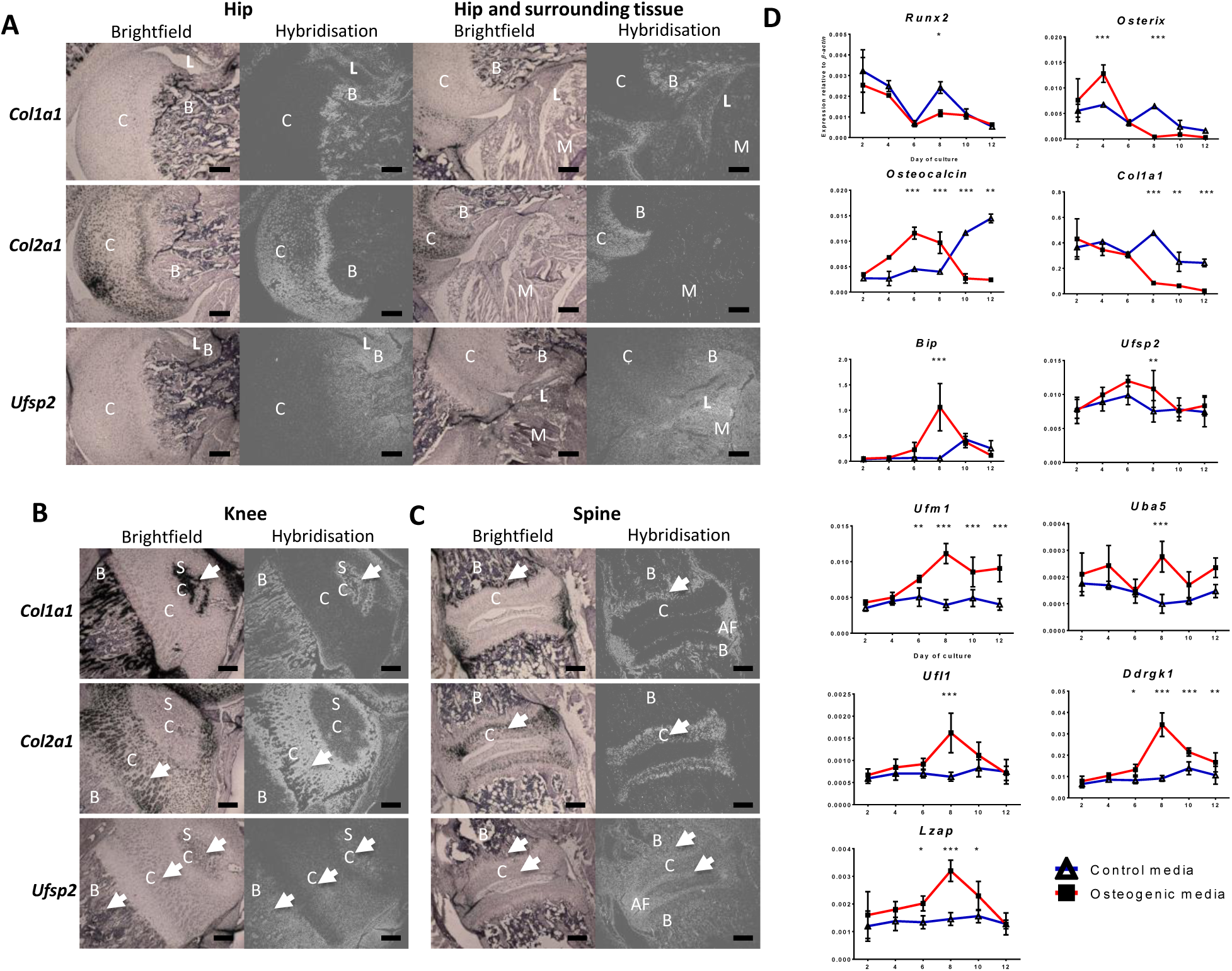
The Ufm1 system components are expressed in mouse joint tissues and upregulated during osteogenic differentiation. ^35^S labelled *Col1a1, Col2a1* and *Ufsp2* RNA probes were hybridised against 10 day old mouse hip (A) knee (B) and spine (C) tissue sections. C – cartilage; B – bone; SC – secondary ossification centre; AF – annulus fibrosus. C – cartilage; B – bone; L – ligament; M – muscle; D. Expression of the Ufm1 pathway during 12 day osteogenic differentiation of 2T3 osteoblasts. Expression was quantified relative to *β-actin*. Values are the mean +/− standard deviation of two independent experiments performed in triplicate. Statistical analysis was performed using 1-way ANOVA with Bonferroni’s post test. * = p<0.05; ** = p<0.01; *** = p<0.001

We performed *in vitro* analysis of the expression of the Ufm1 system during the process of osteogenic differentiation using the 2T3 mouse osteoblast cell line [7]. The mRNA was extracted following 2, 4, 6, 8, 10 and 12 days of culture and gene expression assessed by qPCR. The relative expression of *Runx2, Osterix, Osteocalcin* and *Col1a1* were measured in the 2T3 culture system as the markers of osteogenic differentiation [7]. *Bip* expression was also measured as its expression is known to increase during osteogenic differentiation as well as during ER stress [6]. We found that *Ufsp2, Uba5* (E1), *Ufl1* (E3), *Ddrgk1* and *Lzap* were all significantly upregulated in the induced cultures relative to the control at day 8. *Ufm1* expression started to increase at day 6 to peak at day 8 and remained at this level on days 10 and 12. Thus, in the induced cultures, *Ufsp2, Uba5, Ufl1, Ddrgk1* and *Lzap* upregulation coincided with elevated ER stress as evidenced by upregulation of *Bip*.

### 3.2 Ufm1 system is upregulated in response to ER stress

Based on expression of *Ufsp2* in the mouse joint we used three different cell lines to test the response of the Ufm1 system to ER stress: 2T3 (osteoblasts), C3H 10T½ (embryonic cells that can be differentiated into chondrocytes) and C2C12 (myoblasts). The expression of all the genes tested was upregulated in the DTT treated cultures as compared to untreated controls (Figure 2A). The specificity of the effect of ER stress on the expression of the Ufm1 system in the 2T3 osteoblast cell line was then tested by inducing ER stress with chemical agents acting on different properties of the ER. In addition to DTT we induced ER stress with tunicamycin (Tm) and thapsigargin (Tg). As shown in Figure 2B and C all of Ufm1 system genes tested were upregulated significantly in response to both agents.

**Figure 2.**
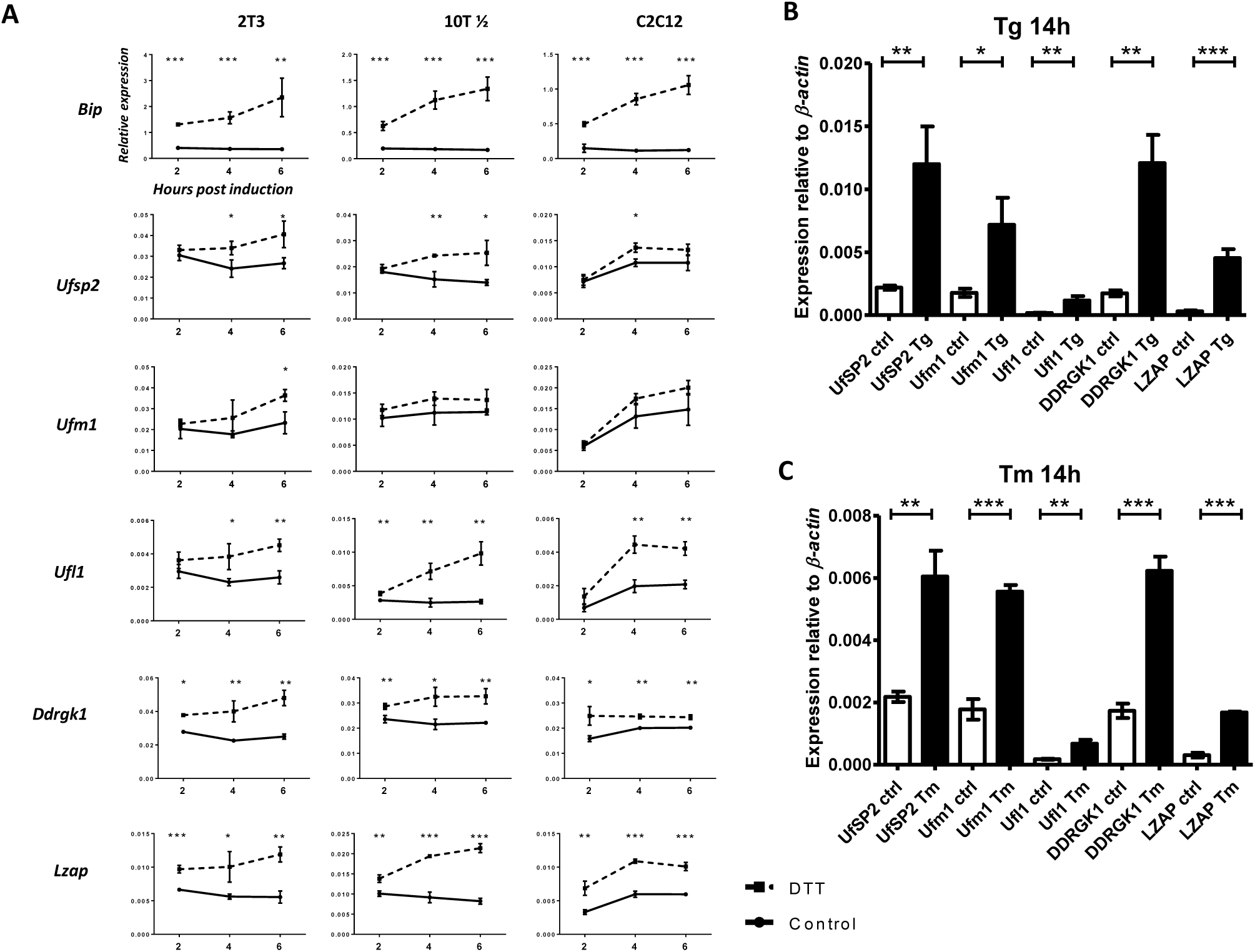
The Ufm1 pathway is upregulated following induction of ER stress. A. ER stress induced in three cell lines (2T3 osteoblasts, C3H 10T½ fibroblasts (10T), and C2C12 myoblasts) by treating the cell cultures with 2mM DTT for 2, 4 and 6 hours. Untreated cultures served as controls. B. and C. ER stress was induced in 2T3 cells by 14 hours of treatment with 2μg/ml tunicamycin (Tm) or 2μM thapsigargin (Tg). Untreated cultures served as controls (ctrl). Gene expression was tested by qPCR. Values are the mean +/− standard deviation of one experiment performed in triplicate. Statistical analysis was performed using the unpaired t test. * = p<0.05; ** = p<0.01; *** = p<0.001.

### 3.3 Genes in the Ufm1 pathway possess Unfolded Protein Response Elements (UPREs) in their promoter regions

Upstream sequences of *Ufsp2, Ufm1, Uba5, Ufc1, Ufl1, Ddrk1, Lzap (Cdk5rap3)* from rat, mouse, human and cattle were aligned. Highly conserved regions in the alignments were analysed for the presence of known regulatory sequences. This analysis revealed the presence of UPREs in the promoter regions of 4 of the 7 known Ufm1 system genes. The UPRE GACGTG sequence potentially bound by the spliced form of XBP1(S), a key transcription factor involved in unfolded protein response [8], was found to be present and highly conserved in the promoter regions of *Ufm1, Uba5, Ufl1* and *Lzap* (Supplementary Figure 2).

To test whether the UPRE sequence played a role in the regulation of *Ufm1, Uba5, Ufl1* and *Lzap* under conditions of ER stress, approximately 1.5 kb of the mouse upstream promoter regions of the above genes were cloned into the pGL3-Basic luciferase vector. The *Ufl1* promoter sequence was not cloned as the sequence failed to amplify after multiple attempts possibly due to low complexity of the genomic region. Mutant versions of the promoter regions (where the UPRE sequence G<u>ACG</u>TG was mutated to G<u>TAA</u>TG) were generated. 2T3 cells were transfected with the promoter-pGL3 vectors and 24 hours post transfection ER stress was induced by addition of Tg. Expression of the luciferase gene driven by the *Uba5* and *Lzap* upstream regions increased by approximately 56% and 21%, respectively following Tg treatment. However, luciferase driven by the *Ufm1* promoter region was noticeably reduced following treatment with Tg. Mutation of the UPRE site significantly decreased the promoter activity of all three constructs. Moreover, treatment with thapsigargin of cells transfected with the mutated promoters did not increase the luciferase expression (Figure 3A).

**Figure 3.**
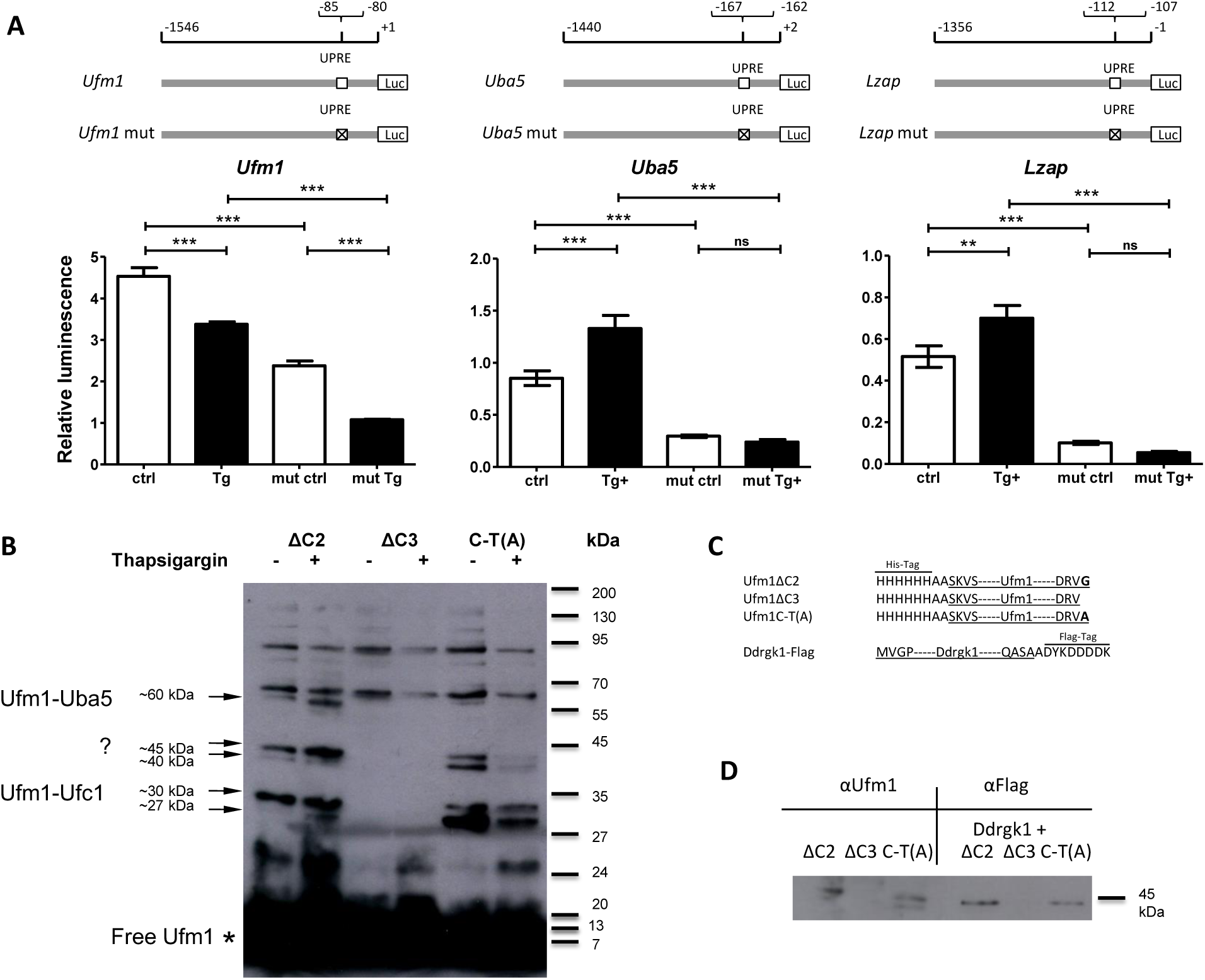
Ufm1 conjugation increases following ER stress in osteoblasts. A. The ruler at the top represents the length of the genomic sequence and position of the UPRE in respect to the ATG codon in mouse genome. UPRE sequences (GACGTG) are marked with open squares and mutated UPRE sequences (GTAATG) are marked with crossed squares. Firefly luciferase expression driven by the native and mutated (mut) *Ufm1, Uba5* and *Lzap* promoter regions under normal (ctrl) and ER stress conditions induced with thapsigargin (Tg) was determined as a ratio of firefly to control *Renilla* luciferase luminescence. Values are the mean +/− standard deviation of one experiment performed in triplicate. Statistical analysis was performed using 1-way ANOVA with Bonferroni’s post-test. * = p<0.05; ** = p<0.01; *** = p<0.001. B. 2T3 osteoblasts were transfected with 3 tagged variants of Ufm1 (ΔC2 – active, ΔC3 – defective and C-T(A) – non-deconjugateble). The proteins were isolated by purification on a Ni-NTA resin from non-stressed cells (-) and cells induced with thapsigargin for 14h (+) and detected by the anti-Ufm1 antibody. Putative Uba5 and Ufc1 bands are marked → as is the possible conjugation target. C. Schematic representation of the Ufm1ΔC2 (activated), Ufm1ΔC3 (conjugation defective) and Ufm1C-T(A) (undergoes permanent conjugation) expression vectors D. Co-purification of overexpressed Ddrgk1 using Ufm1 as bait. 2T3 cells were transfected with 3 tagged variants of Ufm1 only (left), or co-transfected with Flag-tagged Ddrgk1 (right). Proteins were isolated from the cell lysate using His-tag present on Ufm1 and detected by anti-Ufm1 antibody (left) or anti-Flag antibody (right) present on Ddrgk1.

### 3.4 Ufm1 conjugation increases in response to ER stress

2T3 cells were transfected with Ufm1ΔC2 (activated), Ufm1ΔC3 (conjugation defective) and Ufm1C-T(A) (undergoes permanent conjugation) expression vectors (Figure 3C). ER stress was induced with thapsigargin 32 hours post transfection and the cells were harvested 48 hours post transfection (16 hours of induction). Proteins were purified by His-tag present on the overexpressed constructs and detected using anti-Ufm1 antibody (Figure 3B). In the cells transfected with Ufm1ΔC2 there is a visible difference between uninduced and induced cells with the latter having more intensive bands specific to Ufm1 conjugates. In contrast, induction of ER stress had the opposite effect on conjugation or stability of conjugated proteins in the case of the permanent modification with Ufm1C-T(A). The bands at 27, 40 and 45 kDa are less intensive following induction of ER stress and the band at 55kDa (the Ufm1-Uba5 conjugate) is absent. Using mass spectrometry we identified the band at ∼60kDa to be Uba5 and the ∼27-30kDa to be Ufc1 but did not identify the conjugated protein in the ∼40-45kDa band (Figure 3B). Based on literature we suspected that the unidentified band may be Ddrgk1. To confirm this, we co-expressed C-terminally Flag-tagged Ddrgk1 with the His-tagged Ufm1ΔC2, Ufm1ΔC3 and Ufm1C-T(A) expression vectors. The transfected cells were lysed and proteins were purified by the His-tag present on the Ufm1. The purified material was then detected using the anti-Flag antibody which should detect only overexpressed Ddrgk1. The anti-Flag antibody detected a distinct band at a MW of approximately 45 kDa in lanes with proteins purified from Ufm1ΔC2 and Ufm1C-T(A) transfected cultures but no band was present in the conjugation defective Ufm1ΔC3 transfected sample (Figure 3D lanes marked αFlag). The purified Ddrgk1 band corresponded in size to the 45 kDa band detected by the Ufm1 antibody suggesting a Ufm1-Ddrgk1 conjugate (Figure 3D lanes marked αUfm1).

## 3. Discussion

As a first step towards understanding how the mutation in *Ufsp2* leads to the BHD and SEMD phenotype, we determined its expression pattern by radioactive mRNA *in situ* hybridisation against sections of embryonic and postnatal mouse hip joints. *Ufsp2* expression appears to increase postnatally within tissues of the hip joint and it is expressed during secondary centre ossification (as shown in the knee joint). In human development, the secondary centre of ossification of the proximal femoral epiphyses appears at ∼8 months of age and ossification is not complete until 16 years of age. In contrast, in the knee joint, the secondary centre of ossification of the distal femur is present at birth and that of the proximal tibia forms around 1-3 months with ossification being complete by 13-15 years in girls and 15-18 years in boys [9-11]. Thus, early ossification of the proximal femoral head coincides with the time that infants start to walk and hence the hip joint is load bearing. The impact of a delay or alteration in secondary centre ossification (which is seen both in BHD and the SEMD patients [1][2]) may therefore have a greater effect on the hip joint than on the knee joint. The SEMD phenotype also involved other joints and spine however the hip joint was affected the most. Indeed, we have detected expression of *Ufsp2* in the bone, muscles and the secondary ossification centre of the knee and in the bone of the vertebral bodies, and annulus fibrosus and the end plate of intervertebral discs. The difference in the BHD and SEMD phenotype may be due to the specific nature of the mutations or the different genetic backgrounds of the families. Such effects of the *Ufsp2* mutations can however only be explored fully in an *in vivo* model system.

As *Ufsp2* expression observed in the hip joint co-localised with *Col1a1* expression in osteoblasts, patterns of expression of the components of the Ufm1 system were determined relative to markers of osteogenesis in the 2T3 cell line following stimulation with rhBMP-2. Moderate levels of ER stress are known to play an important role in osteogenic differentiation and there is evidence that the Ufm1 pathway may be related to the ER stress that is induced during ischemic heart disease, myositis and type 2 diabetes [12-14]. All components of the Ufm1 system and the *Ddrgk1* and *Lzap* genes were significantly upregulated relative to the control at day 8 which correlates with the expression of *Bip*, a marker of ER stress. Moreover, following treatment with ER stressors (tunicamycin, thapsigargin and DTT) all the Ufm1 system genes tested were significantly upregulated in 2T3 osteoblasts, C3H/10T ½ embryonic fibroblasts and C2C12 myoblasts suggesting it to be a universal mechanism independent of cell type.

To better understand the mechanism of Ufm1 system upregulation during ER stress, the promoter regions of Ufm1 system genes were analysed for presence of known regulatory motifs. This analysis revealed UPRE sequences in the promoter regions of *Ufm1, Uba5, Ufl1* and *Lzap*. During ER stress these sequences are bound by the XBP1 transcription factor which activates transcription of genes essential for dealing with accumulation of unfolded proteins in the ER [8]. Luciferase gene reporter assay performed on ∼1.5 kb mouse promoter regions showed that *Uba5* and *Lzap* were upregulated following treatment with thapsigargin. Mutation of the UPRE sequence showed a decrease in *Luc* gene expression driven by all promoter regions. Moreover, mutation of the *Uba5* and *Lzap* promoter regions prevented the upregulation of *Luc* gene expression in response to thapsigargin treatment. This signifies the importance of the GACGTG sequence in the promoter region for continuous stable expression as well as upregulation of these genes.

Interpretation of the *Ufm1* promoter assay may be more complex as, in addition to the GACGTG sequence located at (−85) to (−80) upstream of the ATG codon which was included in the promoter construct assayed, another similar sequence exists downstream within the second exon at (+108) to (+113) (Supplementary Figure 3). This site was not included in the promoter construct and may be the site that is responsive to ER stress. This requires further investigation. Existence of an XBP1 binding site in the human promoter of *UFM1*was previously reported by Zhang *et al*. [15] at (−67) to (−54) upstream of the *UFM1* start codon. However, the binding sequence that they identified in their paper (AGGGAGCCGTGGA) does not match any known XBP1 consensus binding sequence and this sequence could not be found in the human *UFM1* promoter sequence available in the NCBI database. The mouse *Ufm1* promoter UPRE sequence reported here located at (−85) to (−80) corresponds to the position (−101) to (−96) of the human *UFM1* promoter. In the human *UFM1* promoter this sequence is followed by a further GACGTG sequence (−95) to (−90) not present in other species tested (Supplementary Figure 3). This sequence differs significantly from the putative XBP1 binding site published by Zhang *et al*. (2012) and is more likely to be the actual XBP1 binding site. In support of the data reported here is that the *Ufm1, Uba5, Ufl1* and *Lzap* promoter regions were among genomic fragments pulled down using XBP1 in a chromatin immuno-precipitation study published by Acosta-Alvear *et al*. [8]. The same study established GACGTG as one of the consensus sequences bound by XBP1.

On the protein level treatment of the 2T3 cell line with thapsigargin resulted in increased intensity of the Ufm1 antibody reactive bands corresponding to the conjugation machinery and the putative target Ddrgk1. Interestingly, loss of function mutation of the *DDRGK1* gene was found to result in Shohat-type spondyloepimetaphyseal dysplasia and genetic deletion of in zebrafish and in mice led to disruption of cartilage formation by decreasing Sox9 protein level. In the absence of Ddrgk1 Sox9 becomes highly ubiquitinated which likely leads to proteasomal degradation [4]. It is possible that conjugation of Ufm1 to Ddrgk1 protects it during ER stress and Ddrgk1 in turn stabilizes other proteins like Sox9 however further studies are needed to determine this.

Many chondrodysplasias result from the inability of chondrocytes or osteoblasts to deal with increasing load in the ER, be it due to mutations in the ER machinery or mutations of ECM structural proteins resulting in their misfolding [16]. Many affected by these disorders develop OA as a secondary consequence. In light of the increasing evidence for the importance of ER stress in skeletal development and the findings reported here it is feasible that the link between the *UFSP2* mutation and the BHD phenotype may be related to impaired ER stress responses in chomdrocytes and osteoblasts of the developing hip joint. Conjugation of Ufm1 may regulate stability of key proteins under ER stress conditions during chondrogenic and osteogenic differentiation and the *UFSP2* mutation may disrupt this process. Further studies are however required to determine how the Ufm1 system modulates ER stress responses and how disruption of these processes caused by the UFSP2 mutation leads to skeletal dysplasia.

## Contributors

MD and GAW designed the study and wrote the manuscript. MD performed the experiments. MD and GAW analysed the data.

## Acknowledgements

We would like to thank Dr Gareth Hyde for the Col1a1 and Col2a1 cDNA probes and Dr Chris Watson for the Ufsp2 cDNA probe. We would also like to acknowledge the Biological Mass Spectrometry and the Histology core research facilities at the School of Biological Sciences, University of Manchester.

---

**Supplementary Figure 1.**
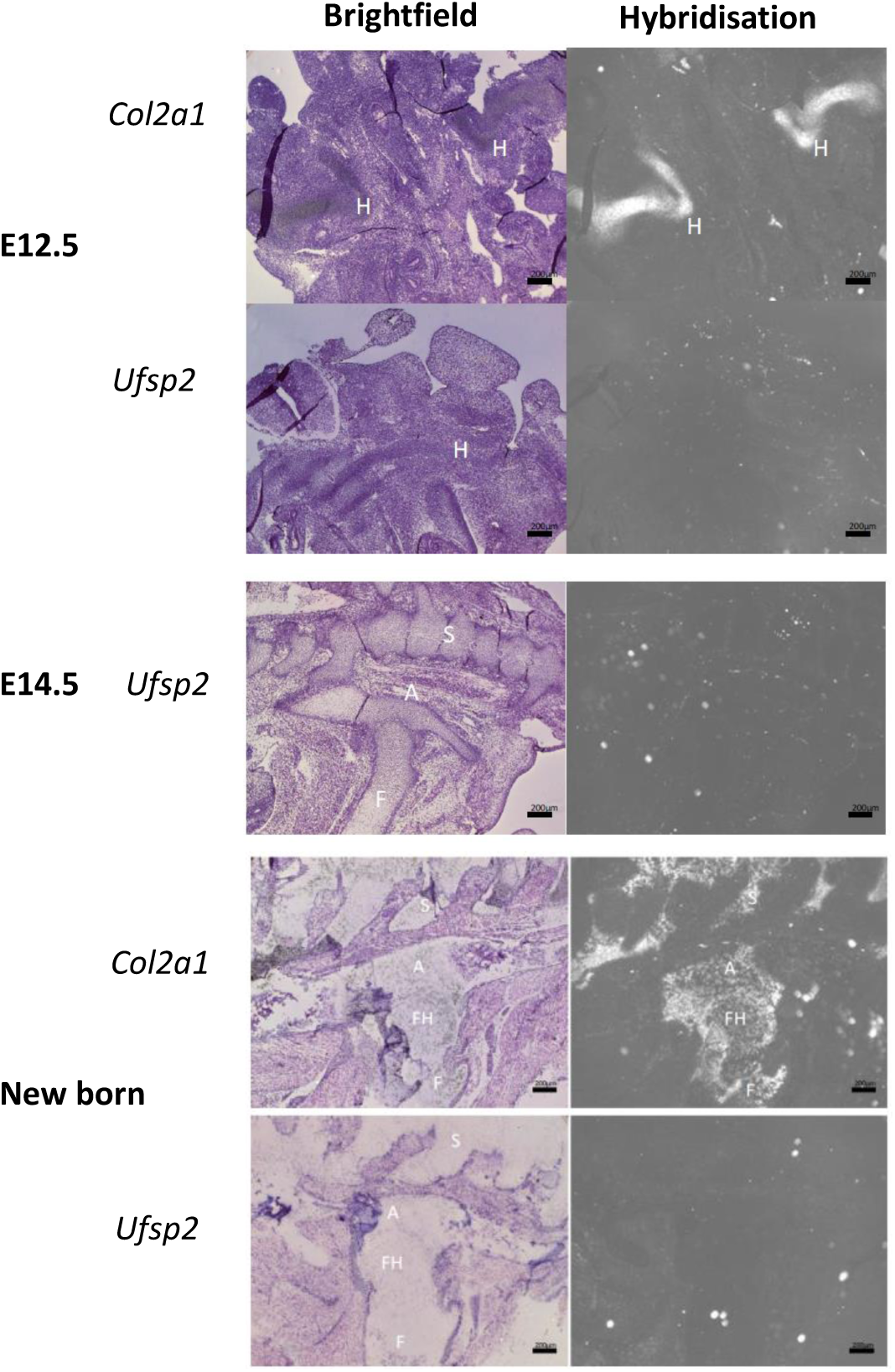
Mouse tissue sections from 12.5 and 14.5 day embryos and new born mice were stained for the expression of *Col2α1* and *Ufsp2*. Panels on the left show H&E staining and panels on the right are images taken in dark-field showing the RNA hybridisation signal. *Col2α1* expression is visible in the cartilage of the presumptive hip joint. No *Ufsp2* expression was detected. H – hip; S – spine; F – femur; FH – femoral head; A - acetabulum. Scale bar 200 μm.

**Supplementary Figure 2.**
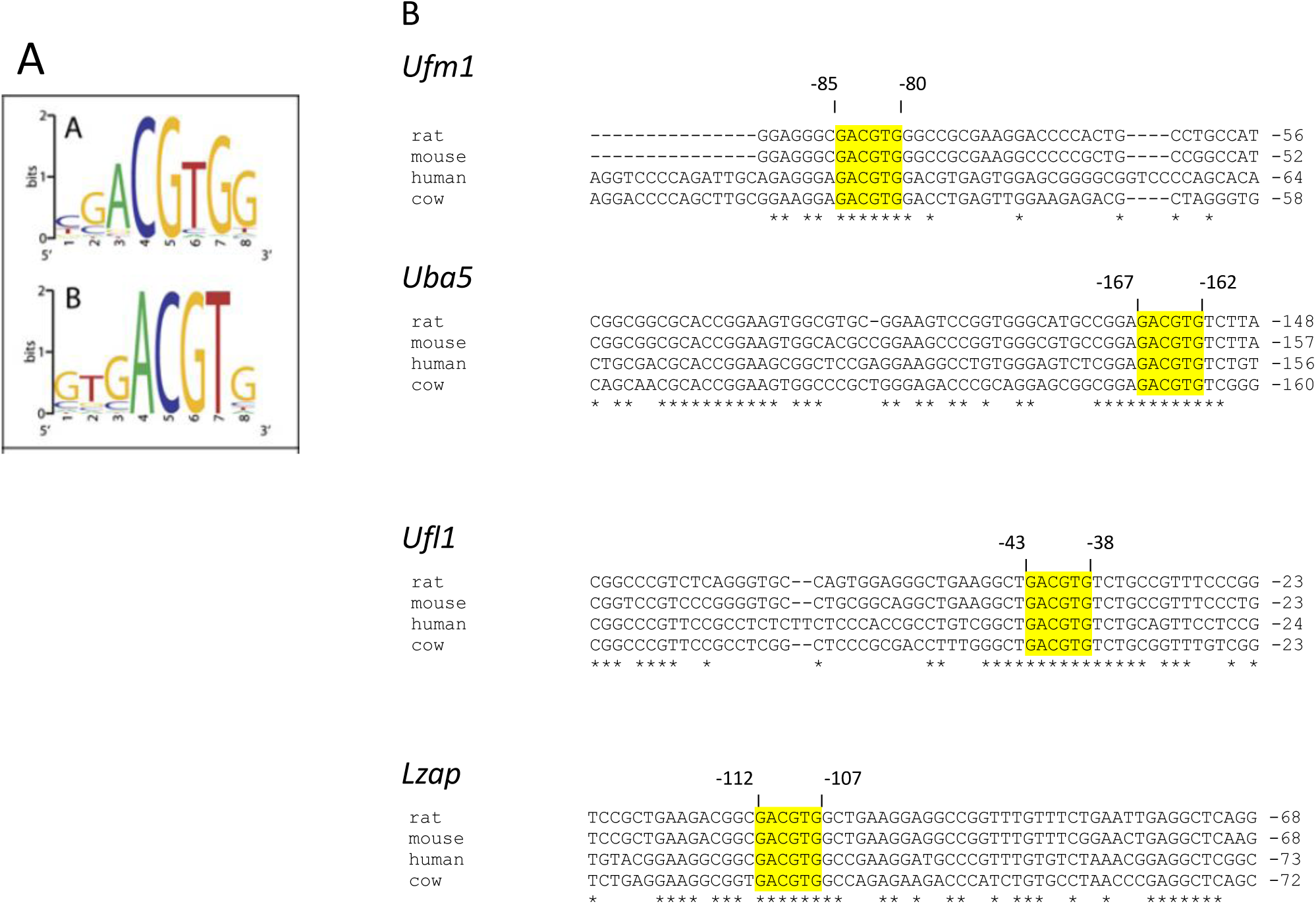
Identification of Unfolded Protein Response Elements (UPRE) in the promoter regions of *Ufm1, Uba5, Ufl1* and *Lzap* genes. A. UPRE consensus sequence bound by the spliced form of Xbp1 derived from chromatin immuno-precipitation studies by Acosta-Alvear *et al* (2007). Size of the letter symbolises frequency of occurrence of this particular residue at this position of the consensus sequence B. Analysis of promoter regions of the *Ufm1* system genes from four different species. The UPRE sequences are highlighted yellow and the numbers denote the position of the UPRE in respect to the ATG codon on the mouse gene. Asterisks denote conserved sequences. UPRE sequences were found in promoter regions of *Ufm1, Uba5, Ufl1* and *Lzap* genes.

**Supplementary Figure 3.**
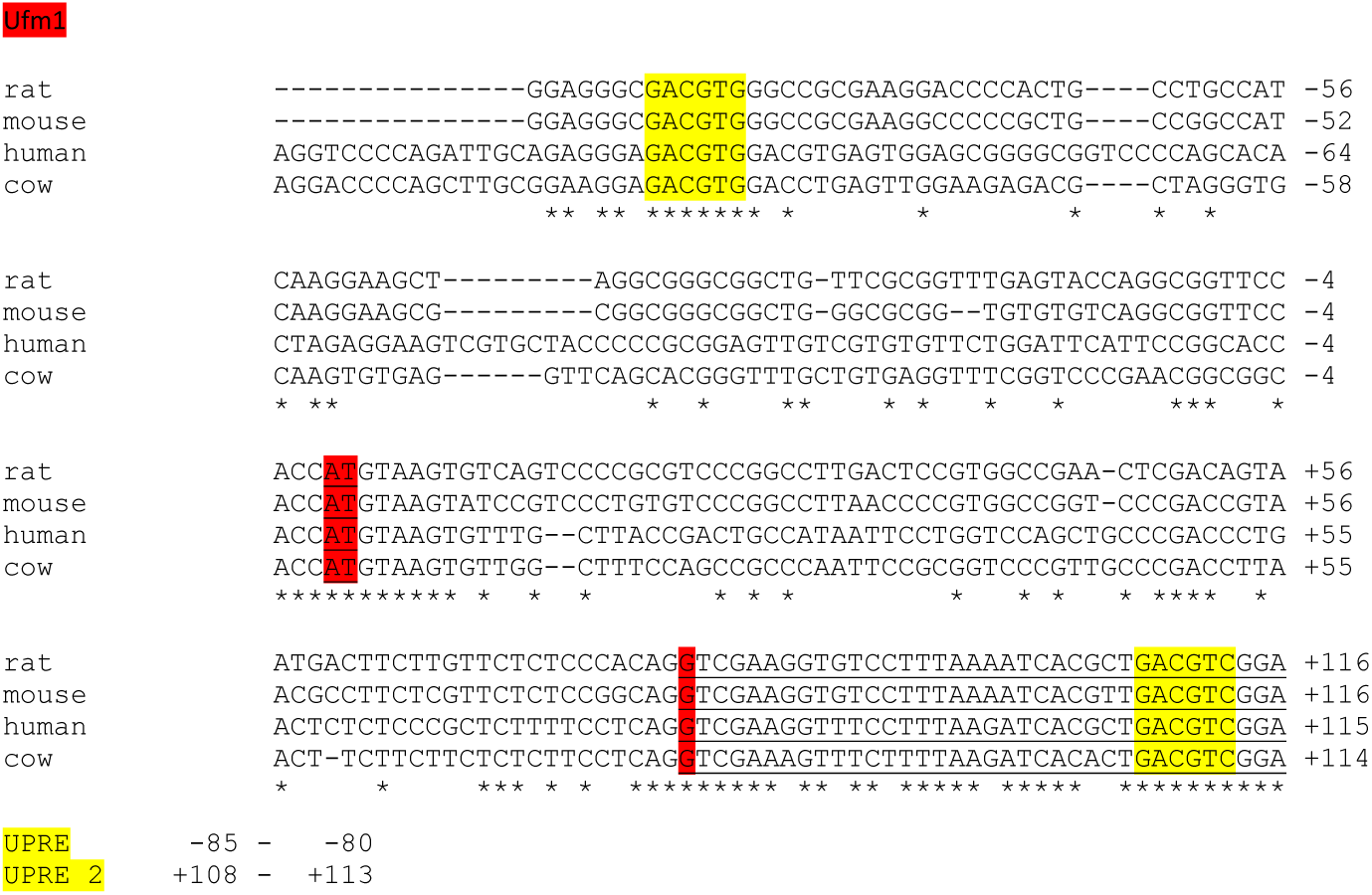
Fragment of the *Ufm1* promoter region including the UPRE sequence. Analysis of promoter region of the *Ufm1* gene from four different species. The UPRE sequences are highlighted yellow. ATG codon is highlighted red. The first and second exon only are underlined. Asterisks denote conserved sequences. Positions of the UPRE sequences in respect to the ATG codon of the mouse gene are given below.

